# Immunotherapy of cytomegalovirus infection by low-dose adoptive transfer of antiviral CD8 T cells relies on substantial post-transfer expansion of central memory cells but not effector-memory cells

**DOI:** 10.1101/2023.08.31.555660

**Authors:** Rafaela Holtappels, Sara Becker, Sara Hamdan, Kirsten Freitag, Jürgen Podlech, Niels A. Lemmermann, Matthias J. Reddehase

## Abstract

Cytomegaloviruses (CMVs) are host species-specific in their replication. It is a hallmark of all CMVs that productive primary infection is controlled by concerted innate and adaptive immune responses in the immunocompetent host. As a result, the infection usually passes without overt clinical symptoms and develops into latent infection, referred to as ‘latency’. During latency, the virus is maintained in a non-replicative state from which it can reactivate to productive infection under conditions of waning immune surveillance. In contrast, infection of an immunocompromised host causes CMV disease with viral multiple-organ histopathology resulting in organ failure. Primary or reactivated CMV infection of hematopoietic cell transplantation (HCT) recipients in a “window of risk” between therapeutic hematoablative leukemia therapy and immune system reconstitution remains a clinical challenge. Studies in the mouse model of experimental HCT and infection with murine CMV (mCMV), followed by clinical trials in HCT patients with human CMV (hCMV) reactivation, have revealed a protective function of virus-specific CD8 T cells upon adoptive cell transfer (AT). Memory CD8 T cells derived from latently infected hosts are a favored source for immunotherapy by AT. Strikingly low numbers of these cells were found to prevent CMV disease, suggesting either an immediate effector function of few transferred cells or a clonal expansion generating high numbers of effector cells. In the murine model, the memory population consists of resting central memory T cells (TCM), as well as of conventional effector-memory T cells (cTEM) and inflationary effector-memory T cells (iTEM). iTEM, increase in numbers over time in the latently infected host, a phenomenon known as ‘memory inflation’ (MI). They thus appeared to be a promising source for use in immunotherapy. However, we show here that iTEM contribute little to the control of infection after AT, which rests almost exclusively on a superior proliferation potential of TCM.

**Author Summary:** Immunotherapy of reactivated cytomegalovirus infection in immunocompromised HCT recipients by adoptive transfer (AT) of antiviral CD8 T cells is the last resort to fight virus variants that have acquired resistance to standard antiviral drugs. Provision of cell numbers high enough for clearance of productive infection remains a logistical limitation for AT to become clinical routine. Although use of donor memory CD8 T cells has become the standard in clinical AT, little is known about the relative antiviral efficacies of memory CD8 T-cell activation subsets, such as central memory cells (TCM) and different populations of effector-memory cells (TEM). A reliable quantitative comparison of the antiviral efficacies of memory CD8 T-cell subsets is precluded in clinical investigation, because independent cohorts of AT donors and AT recipients unavoidably differ in many genetical, immunological, and virological variables. Therefore, this is a question for which a preclinical animal model is predestined. We show here in the well-established mouse model of low-dose AT that CMV infection is by far most efficiently controlled by virus-specific TCM, based on a superior potential to proliferate even in extra-lymphoid tissue to prevent virus spread. For clinical AT, our data provide an argument to favor transfer of sorted TCM rather than TEM.

## Introduction

The clinical relevance of human cytomegalovirus (hCMV), the prototype member of the β-subfamily of the herpes virus family [1], results from severe and often lethal CMV disease that it causes under conditions of compromised immunity [2–4]. A concern with significant health system impact are birth defects resulting from congenital infection of immunologically immature fetuses, a syndrome historically known as cytomegalic inclusion disease (CID) (for overviews, see [5,6]). Risk groups of medical and logistic challenge in transplantation centers worldwide are iatrogenically immunocompromised recipients of hematopoietic cell transplantation (HCT) [7] and of solid organ transplantation (SOT) [8,9]. In those patients, hCMV reactivation within a latently infected transplant or within latently infected organs of the recipient causes graft failure and multiple organ disease, with interstitial pneumonia representing the most threatening clinical manifestation specifically in HCT (for clinical reviews, see [10–13].

As CMVs are host-species specific in their replication based on host range determinants [14–16], understanding of the underlying mechanisms of CMV disease and immune control of infection comes at its limits when research is restricted to clinical investigation. In particular, human genetics cannot be manipulated, and hCMV mutants cannot be used experimentally for designed *in vivo* studies aimed at identifying the roles of host and viral genes involved in pathogenesis and immunity. Of all animal models of CMV disease and infection control, infection of the mouse with murine CMV (mCMV) is the most advanced with respect to host genetics [17]. As hCMV and mCMV differ genetically, each not just containing homologous genes with related functions but also ‘private genes’ co-evolved with and thereby adapted to the respective host species [1,18–20], the results for mCMV cannot be translated par for par to hCMV. Nonetheless, the mouse model has identified basic principles common to all CMVs [21]. As a paramount example for successful clinical translation, mouse models of experimental syngeneic and allogeneic HCT [22–26], for overviews see [27,28], and of control of infection by adoptive transfer (AT) of virus-specific CD8 T cells [29–33] were of predictive value for the management of human infection. Specifically, results from mouse models have paved the way to ‘pre-emptive immunotherapy’ of hCMV infection in HCT recipients. This means prevention of CMV disease by transfer of virus-specific CD8 T cells as soon as virus reactivation is detected in the routine follow-up screening long before clinical diagnosis of CMV disease manifestations [34–38]. Although AT in a clinical scale is still logistically demanding, it is the last resort to fight infection by virus variants that have acquired resistance to standard antiviral drugs [39–45].

The source of CD8 T cells for clinical immunotherapy are mostly latently infected but otherwise healthy donors who bear virus-specific memory CD8 T cells resulting from previous hCMV infection. Ideally, the CD8 T-cell donor is the one who has been MHC class-I matched for the HCT, so that the viral epitope-specificity of the memory CD8 T cells matches with the epitopes presented by the MHC class-I molecules of the combined HCT and AT recipient. Studies conducted on mice ([46–48], for a review see [49]), as well as clinical trials [37,38,50], have consistently demonstrated that memory CD8 T cells are more effective in immunotherapy when compared to terminally-differentiated effector CD8 T cells of *in vitro*-propagated cytolytic T cell lines (CTLL) of identical epitope-specificity. Accordingly, the previous approach of expanding CD8 T cells in cell culture to reach sufficiently high cell numbers for AT [35,51] is no longer pursued. Suspected reasons were loss of functional avidity in the recognition of presented antigenic peptides or a loss of *in vivo* homing properties as a result of selection during expansion in cell culture. In retrospect, our data presented here rather suggest that loss of proliferative capacity by terminal differentiation to effector cells explains the low per-cell antiviral efficacy of CTLL compared to memory CD8 T cells capable of expanding within the AT recipient.

In latently infected mice, the memory CD8 T-cell population consists of three major subpopulations, all expressing CD44 in distinction to naïve, unprimed CD8 T cells: central memory T cells (TCM) characterized by the cell surface phenotype CD62L^+^KLRG1^-^, as well as conventional T effector-memory cells (cTEM) and inflationary effector-memory cells (iTEM) with the cell surface phenotypes CD62L^-^KLRG1^-^ and CD62L^-^KLRG1^+^, respectively [52]. iTEM were previously referred to as short-lived effector cells (SLEC) [53] but were then shown to differ distinctively from terminally-differentiated effector T cells (TEC) by dependence on IL-15 [54]. After high-dose systemic but not after low-dose local infection [52], the iTEM population reflects a phenomenon named “memory inflation” (MI), as numbers of iTEM increase almost steadily over time during latent mCMV infection (for reviews, see [55–58]). It is current view that the latent viral genome is not transcriptionally silent but that sporadic episodes of limited viral gene expression [59–61], proposed to take place in latently infected endothelial cells [61–63] and likely also in latently infected PDGFRα^+^ fibroblasts [64], lead to the presentation of antigenic peptides that drive MI [55].

Here we have sorted TCM, cTEM, and iTEM for AT into immunocompromised and infected recipients to directly quantify their individual contributions to the control of infection. Notably, prevention of viral spread and pathogenesis was based almost entirely on TCM, and correlated with the proliferative capacity that was highest for TCM and lowest for iTEM. In conclusion, efficient expansion of antiviral TCM within the host is crucial for control of infection and prevention of CMV disease upon pre-emptive immunotherapy by AT.

## Results

### Adoptive transfer of remarkably low numbers of virus-specific memory CD8 T cells controls virus replication at multiple organ sites

It is long established in the murine model system, as well as by clinical trials, that lethal CMV infection of the immunocompromised host can be prevented by AT of CMV-specific CD8 T cells, provided that the protective cells are administered early after infection (see the Introduction). This is the basis for clinical preemptive immunotherapy of hCMV reactivation in HCT recipients upon first detection in a sensitive follow-up screening. A favored source of CMV-specific CD8 T cells are memory cells collected from a latently infected but otherwise healthy donor, ideally the MHC-matched HCT donor. This ensures an optimized fit in viral antigenic peptides presented by infected cells in tissues of the recipient and recognized by the CD8 T cells derived from the donor.

The term ‘memory cells’ is here used collectively for a mixture of antigen-experienced cells in different stages of differentiation and activation, comprising resting TCM as well as activated cTEM and iTEM [52]. Our study aimed at quantitating the individual contributions of these memory cell subsets to the control of CMV infection in the well-established mouse model.

In order to calibrate the system, we first studied the unseparated memory cell population. Total CD8 cells derived from the spleen of latently infected BALB/c mice were enriched by positive immunomagnetic cell sorting (Fig 1A). Cytofluorometric analysis revealed presence of the three main memory-cell populations defined by the expression of CD62L and KLRG1, that is, the TCM, cTEM, and iTEM. Almost all cells of the memory population expressed CD44, distinguishing them from CD44^-^ naïve cells that have not yet encountered antigen and represent the minority of peripheral CD8 T cells (S1 Fig). As noted in the important work by Welsh and Selin, ‘no one is naïve’ [65], so that memory cells of numerous antigen specificities make up the memory cell population. Trivially, most memory CD8 T cells are not specific for the virus under investigation but collectively reflect all preceding antigen encounters in the past life. The mCMV-specific fraction of memory cells was determined by an IFNγ-based ELISpot-Assay covering all response-relevant CD8 T-cell epitopes of mCMV in the *H-2^d^* haplotype (for a list of peptide sequences and the presenting MHC class-I molecules, see [49]) (Fig 1B). In accordance with previous data [24,66], memory CD8 T cells specific for the epitopes IE1 and m164 dominated the response, and cells specific for the tested panel of epitopes added up to ∼0,9 % of the population. Other already identified epitopes, which were not included in the analysis, are known to generally contribute little to the overall CD8 T-cell response, and previous work, using a viral genome-wide open reading frame (ORF) expression library [67], did not reveal the existence of unidentified CD8 T-cell epitopes that would make a notable contribution to the memory response [68]. It is therefore reasonable to conservatively extrapolate that mCMV epitope-specific memory cells accounted for not more than ∼1 % of the CD8 T-cell population. Given the universe of antigens, this is nevertheless a remarkable allocation of the memory CD8 T-cell pool to a single viral pathogen.

**Fig 1.**
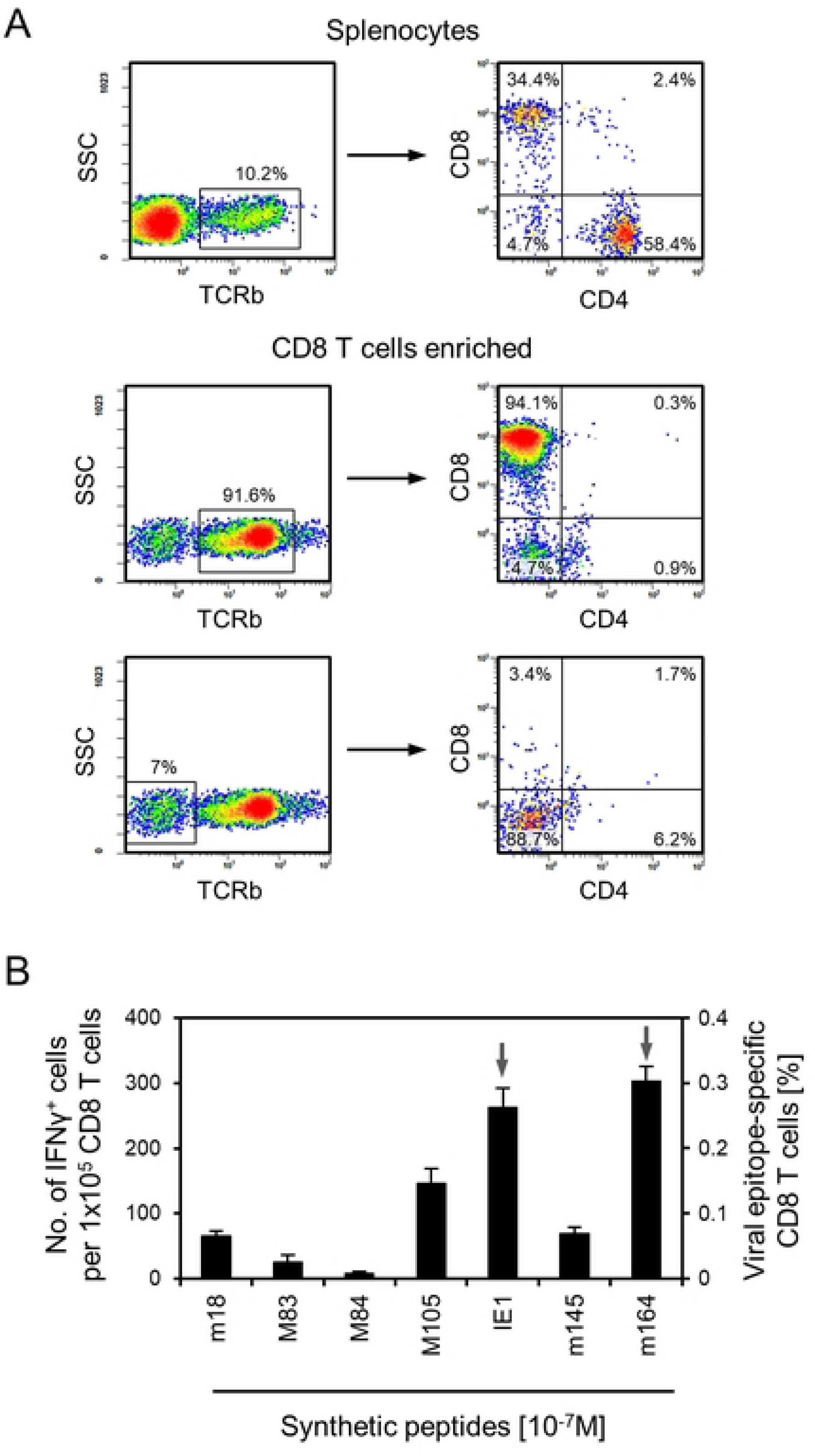
Characterization of the donor memory CD8 T cells used for AT. Splenocytes were isolated from latently infected BALB/c mice, and CD8^+^ cells were enriched by positive immunomagnetic cell sorting. (A) CFM analyses documenting the proportions of CD8 and CD4 T cells, within the gate on T cells defined by TCR (TCR β-chain) expression, before and after immunomagnetic enrichment of CD8^+^ cells (upper and center panel, respectively). Gating on TCR-negative cells in the CD8^+^ cell-enriched splenocyte population reveals that CD8^+^ dendritic cells (DC) are numerically negligible (lower panel). Shown are color-coded 2D fluorescence density plots with red and blue color representing highest and lowest cell numbers, respectively. (B) Frequencies of memory CD8 T cells specific for the viral epitopes indicated. Bars represent the frequencies of cells stimulated by viral antigenic peptides to secrete IFNγ in an ELISpot assay, determined by intercept-free linear regression analysis. Error bars represent the 95% confidence intervals. Arrows highlight the frequencies of the two immunodominant CD8 T-cell specificities IE1 and m164.

The thus characterized cells were then used as donor cells for syngeneic AT into immunocompromised BALB/c recipients (for a protocol scheme, see Fig 2A). Extensive viral replication in vital organs at read-out time would lead to lethal CMV disease with tissue lesions if left untreated. Notably, as few as 1,000 transferred cells, containing just ∼10 viral epitope-specific cells (Fig 1B), significantly reduced viral replication in three organs tested, namely in spleen, lungs, and liver. Almost clearance of the infection was achieved by AT of 100,000 cells, corresponding to ∼1,000 viral epitope-specific cells (Fig 2B).

**Fig 2.**
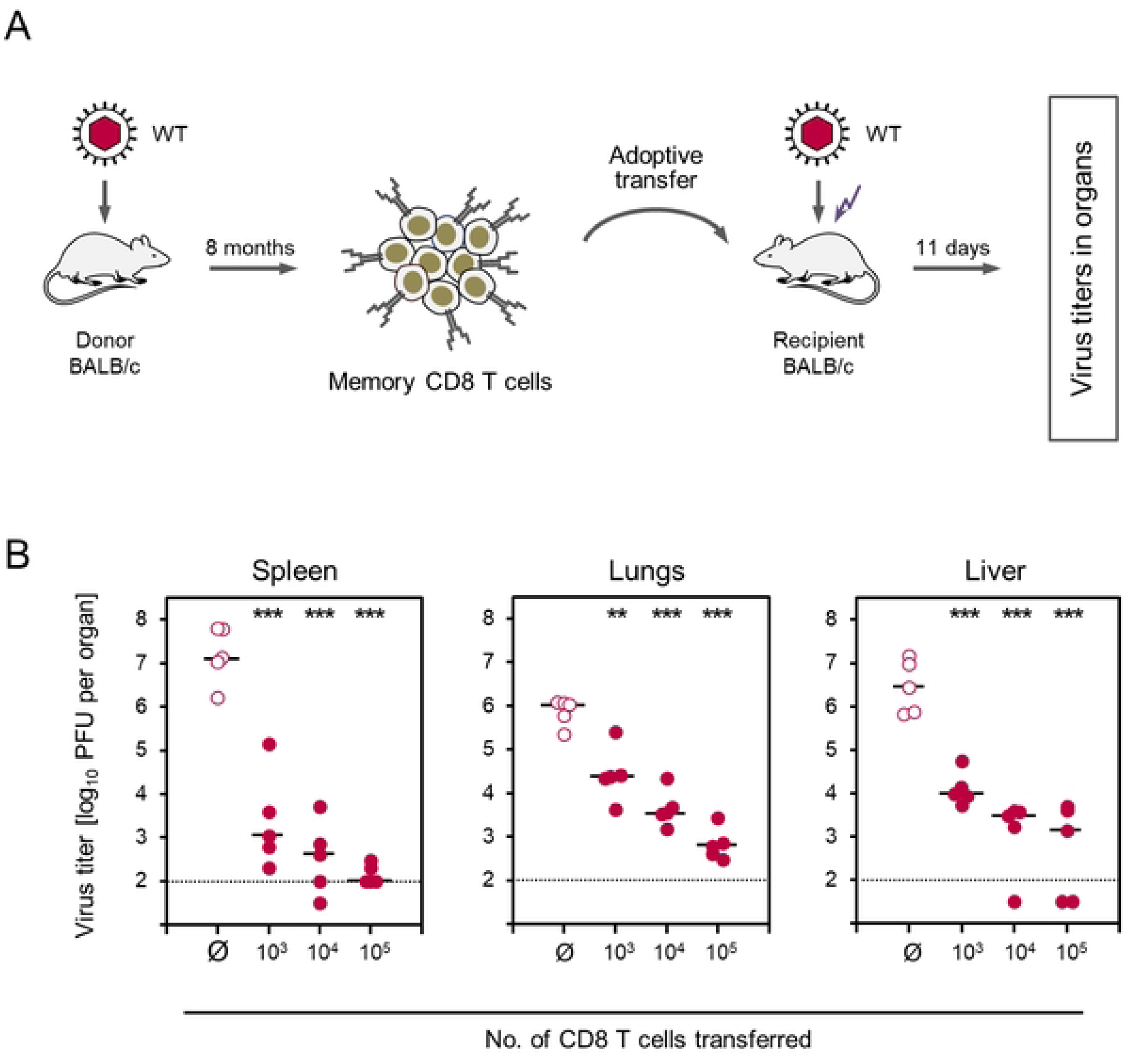
Control of infection by AT of memory CD8 T cells. (A) Sketch of the experimental protocol. Total memory CD8 T cells, which include viral-epitope specific cells in frequencies shown in Fig 1B, were transferred into BALB/c recipient mice that were immunocompromised by γ-irradiation (flash symbol) and infected with mCMV-WT (WT). (B) Virus titers in spleen, lungs, and liver determined on day 11 after AT of graded cell numbers indicated. Ø, no AT performed. Symbols represent data from individual mice. Median values are marked. Dotted lines represent the detection limit of the virus plaque assay. (PFU) plaque-forming units. Asterisk-coded statistical significance levels for differences between the AT groups and the no-AT control group (Ø): P < (**) 0.01 and (***) 0.001.

### Control of infection is mediated by viral epitope-specific memory CD8 T cells

We have previously reported a strategy to test for viral epitope-specificity of CD8 T-cell priming, target cell recognition, and protection against *in vivo* virus replication in AT models by site-directed virus mutagenesis of the C-terminal amino acid residue that anchors an antigenic peptide to the presenting MHC class-I molecule [69]. This strategy was first applied to the deletion of IE1 peptide antigenicity and immunogenicity in mCMV mutant IE1-L176A, in which the MHC class-I (L^d^) anchor residue Leu at the C-terminal position of the antigenic IE1 peptide was replaced with Ala [60]. More recently, this strategy was employed to show viral epitope-specificity of antiviral protection upon AT in a mouse model of ‘humanized antigen presentation” using TCR-transduced murine or human CD8^+^ CTLL specific for an antigenic peptide of hCMV presented on tissue cells of HLA-A2 transgenic mice [31]. Notably, AT of these CTLL, both murine and human CTLL, into recipients infected with recombinant mCMV expressing the authentic antigenic peptide led to tissue infiltration by CD8 T cells, associated with the formation of ‘nodular inflammatory foci (NIF)’ to which infection is confined and eventually cleared. In contrast, when AT recipients were infected with recombinant mCMV expressing the C-terminal Ala-variant of the peptide, infiltrates were missing almost completely and the virus spread unhindered with consequent viral histopathological lesions [31].

Here we tested if epitope-specificity of antiviral protection also applies to *ex vivo* sorted IE1 epitope-specific memory cells that differ from CTLL in activation stage, proliferative capacity, and tissue homing properties. For this, immunocompromised AT recipient mice were infected either with the mCMV mutant IE1-L176A, in which presentation of the IE1 epitope is largely reduced, or with the corresponding revertant IE1-A176L (Fig 3). When no AT was performed, two-color immunohistochemical (2C-IHC) analysis of representative liver tissue sections stained for the CD8 molecule and the viral IE1 protein revealed absence of CD8 T cells and comparable tissue spread of both viruses. Viral spread becomes visible as extended tissue areas with infected IE1^+^ liver cells, which are predominantly hepatocytes (iHc) and endothelial cells (iEC) [70,71]. When AT was performed, CD8 T cells infiltrating liver tissue infected with the mutant virus were found in proximity to infected cells, but controlled infection inefficiently. In contrast, after infection with the revertant virus, CD8 T cells clustered more densely around infected cells to form NIF, thereby confining and eventually resolving the infection (Fig 3A). Quantitation of liver tissue-infiltrating CD8 T cells and of infected liver cells in the respective groups of mice confirmed a significantly better control of the revertant virus expressing the authentic antigenic IE1 peptide that binds with high affinity to the presenting MHC class-I molecule (Fig 3B).

**Fig 3.**
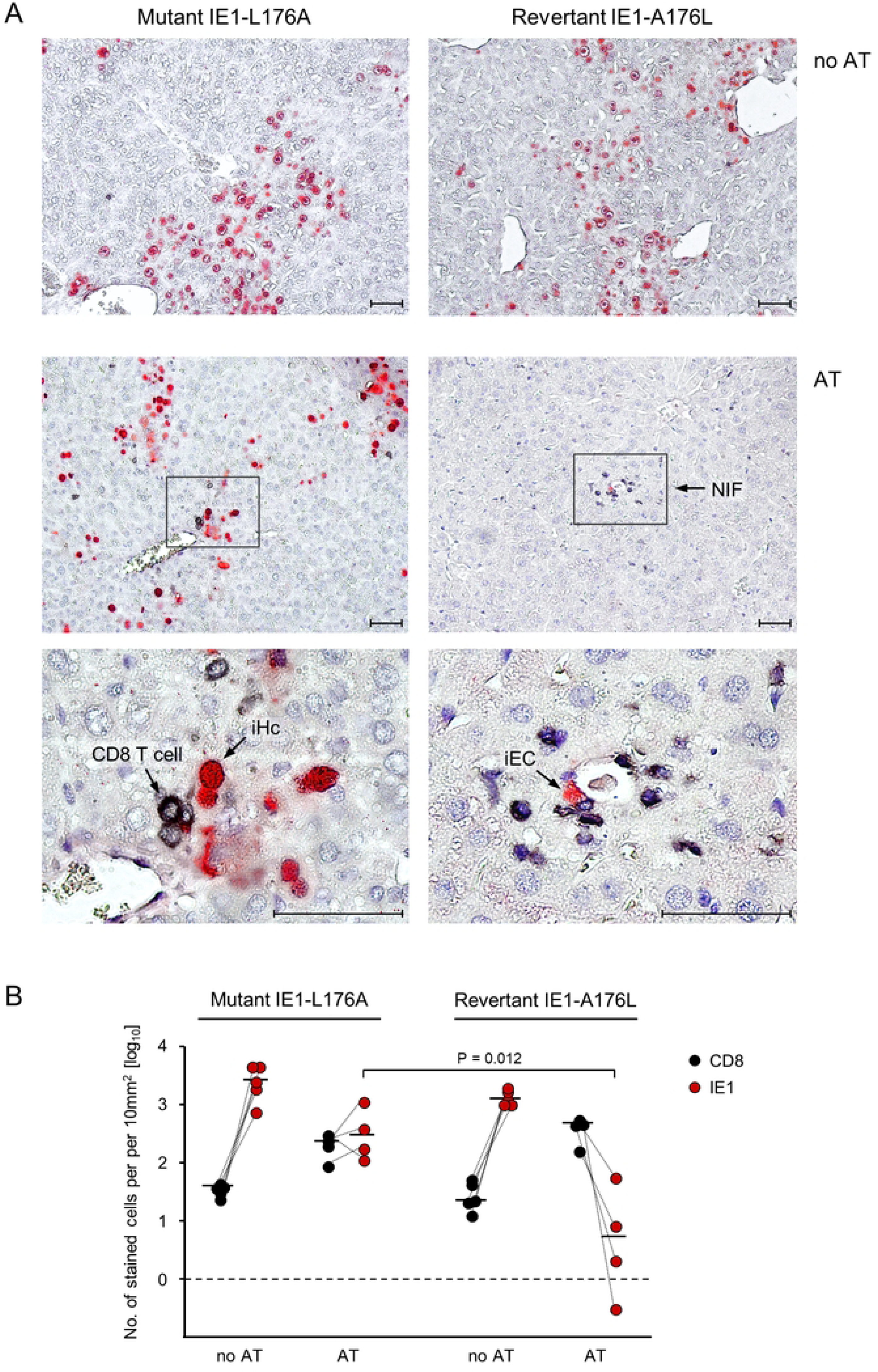
Viral epitope-specificity of antiviral control. AT was performed with 10^4^ IE1-epitope specific memory CD8 T cells isolated from splenocytes of latently-infected BALB/c donor mice by negative immunomagnetic pre-enrichment of CD8^+^ cells followed by fluorescence-based sorting of cells expressing IE1 epitope-specific TCRs. AT recipients were infected either with the IE1 epitope mutant virus mCMV-IE1-L176A or the corresponding revertant virus mCMV-IE1-A176L. Analyses were performed on day 11 after AT and infection. (A) 2C-IHC images of liver tissue sections. (Left panels) AT recipients infected with mCMV-IE1-L176A. (Right panels) AT recipients infected with mCMV-IE1-A176L. Infected liver cells are identified by red staining of the intra-nuclear viral protein IE1. (iHc) infected hepatocyte. (iEC) infected capillary endothelial cell. Tissue-infiltrating IE1-specific CD8 T cells are identified by black staining of the CD8a molecule. Light hematoxylin counterstaining reveals the context of liver tissue. (Upper panel, no AT) Unhindered intra-tissue spread of both viruses. (Center and lower panels, AT) The center panel provides a low-magnification overview, showing the confinement of infection by formation of nodular inflammatory foci (NIF) only after infection with revertant virus mCMV-IE1-A176L expressing the antigenic IE1 peptide on infected cells. Frames demarcate regions of interest resolved to greater detail in the lower panel images. Bar markers represent 50µm. (B) Quantitation of infected IE1^+^ liver cells (red dots) and tissue-infiltrating CD8 T cells (black dots) in representative 10-mm^2^ liver tissue section areas. Symbols represent cell counts for AT recipients tested individually. Linked data are connected by dotted lines, median values are marked. The dashed line indicates the detection limit of the assay.

Some tissue infiltration by CD8 T cells and partial control of mutant virus IE1-L176A might be explained by low-affinity MHC class-I binding of the L176A peptide [60,72], sufficient for presenting the TCR contact site 170-HFMPT-174 [72]. Although the L176A mutation is predicted to also reduce the probability of proteasomal C-terminal cleavage [73–75], trace amounts of the mutated peptide may bind to the presenting MHC class-I molecule L^d^, catalyzed in the peptide loading complex [76,77]. Such a limited presentation may suffice for sensitization of a fraction of the polyclonal IE1-specific memory CD8 T cells that possess TCRs of particularly high avidity.

### Differential antiviral efficacy of memory CD8 T-cell subsets

Data so far essentially confirmed recent findings reported for AT performed with total memory CD8 T cells [78]. Here we expanded on this study by investigating the differential contributions of memory CD8 T-cell subsets representing activation stages defined by the expression of the cell surface markers CD62L and KLRG1: T central memory cells (TCM) display the cell surface phenotype CD62L^+^KLRG1^-^ and are distinct from naïve T cells by the expression of CD44. T effector-memory cells (TEM) lack expression of CD62L and split into conventional TEM (cTEM) and inflationary TEM (iTEM), distinguished by absence or presence of KLRG1 cell surface expression, respectively (S2B Fig).

We were particularly interested in quantitating the antiviral potency of the CD62L^-^ KLRG1^+^ iTEM, which are highly activated due to more recent restimulation by the cognate viral antigenic peptide. This is expressed and presented in a stochastic manner during mCMV latency as a result of intermittent transcriptional gene de-silencing events [61]. They expand and thus accumulate over time based on repetitive restimulation, a phenomenon known as MI. Based on their original classification as ‘short-lived effector cells’ (SLEC) [53], iTEM are supposed to interfere with virus replication almost instantly after AT already at the site of infection and prevent virus dissemination to organ sites of CMV disease. In contrast, TCM must first see the cognate epitope for clonal expansion and maturation to effector cells. They therefore might come too late for preventing virus colonization of host tissues and might rather interfere with later intra-tissue virus spread. In fact, this was recently found to be the case for unseparated total memory CD8 T cells [78], but in that earlier study the quantitative subset composition of the memory cell pool and possible differential contributions of memory CD8 T-cell subsets, temporally and spatially, were not addressed. Thus, the previous findings may have reflected the properties of the majority subset. As we know now, the TCM represent the majority subset (Fig 4B, S2B Fig).

**Fig 4.**
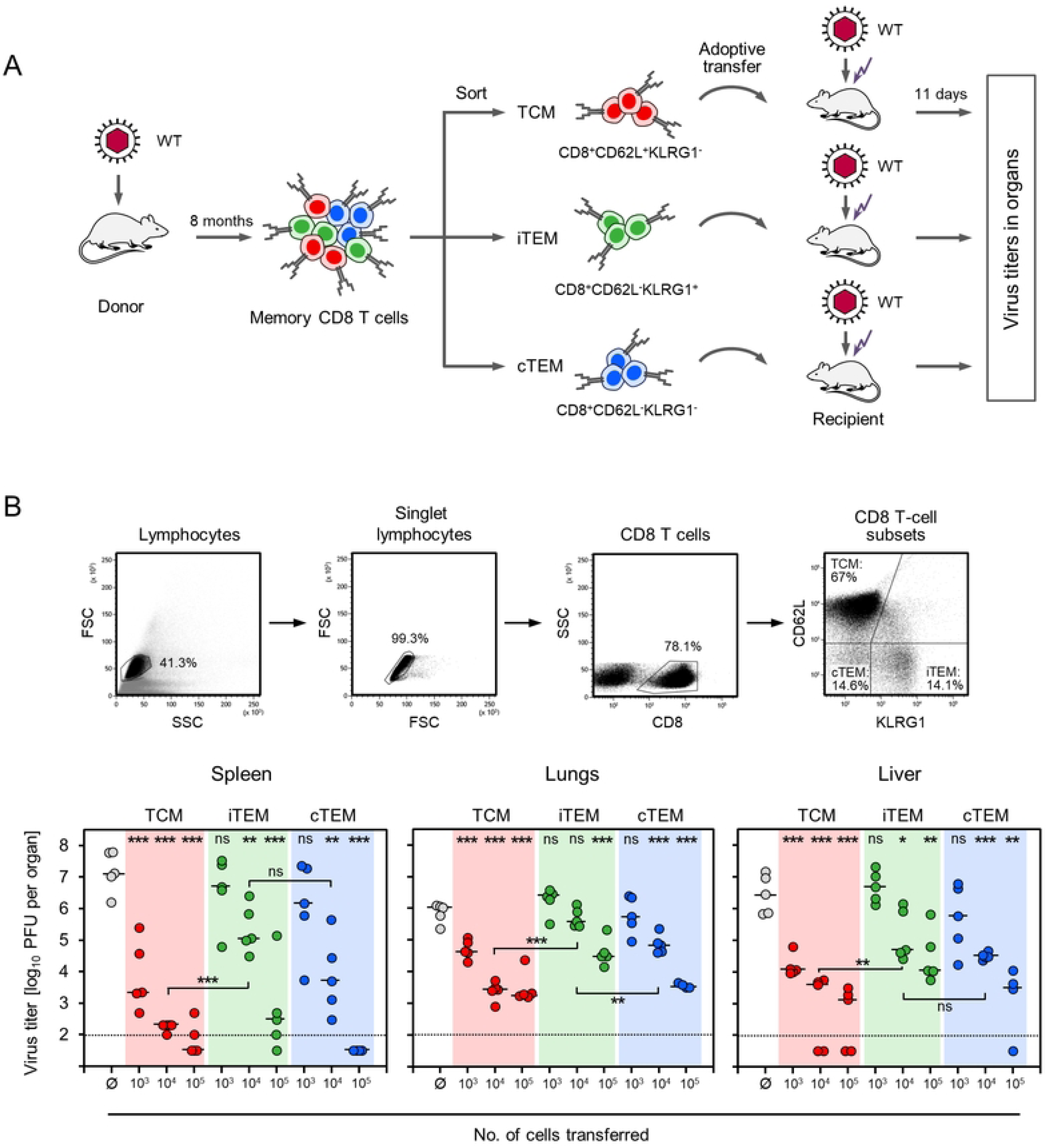
Control of infection by AT of memory CD8 T-cell subsets. (A) Sketch of the experimental protocol. (WT) mCMV-WT. (TCM) T central memory cells. (iTEM) inflationary T effector-memory cells. (cTEM) conventional T effector-memory cells. For AT recipients, see the Legend to Fig 2. (B, upper panel) Gating strategy for the cell sorting. Shown are 2D-dot plots with progressing gates set on the cells indicated. (FSC) forward scatter. (SSC) sideward scatter. (B, lower panel) Control of productive infection in organs of AT recipients by sort-purified donor-derived memory CD8 T-cell subsets. AT was performed with graded donor cell numbers. Ø, no AT performed. Symbols represent individual mice with median values indicated. Data for transferred CD8 T-cell subsets are color-coded as defined in (A). Dotted lines represent the detection limit of the virus plaque assay. (PFU) plaque-forming units. Asterisk-coded statistical significance levels for differences between the AT groups and the no-AT control group (Ø): P < (*) 0.05, (**) 0.01, and (***) 0.001. (n.s.) not significant.

For evaluating the contributions of the three subsets separately and normalized to a per-cell basis, they were purified by fluorescence-based cell sorting and tested for their individual antiviral efficacies upon AT into infected, immunocompromised recipient mice (S2 Fig, pilot experiment; Fig 4, reproduction in a second experiment with an extended range of cell numbers transferred). The two independent experiments were consistent in showing that TCM were the most potent memory cell subset in controlling virus replication in host organs, whereas iTEM were the least potent. Specifically, when compared to the control group of no AT, 1,000 TCM significantly reduced virus replication in all three organs tested, whereas iTEM and cTEM both failed (Fig 4). With increasing cell numbers, control of infection by iTEM and cTEM was increasingly improved, with a tendency to the favor of cTEM over iTEM. Only upon transfer of 100,000 cells, all three subsets reduced virus replication with statistical significance in all organs tested (Fig 3). In conclusion, the ranking in antiviral efficacy was TCM >> cTEM > iTEM.

### The superior antiviral activity of TCM is not explained by higher numbers of functional viral epitope-specific cells

As protection is mediated by epitope-specific CD8 T cells (Fig 3), differential antiviral activity of sorted memory CD8 T-cell subsets might simply reflect differences in the specificity composition of the cell pools. We focused our analysis on the frequencies of IE1- and m164-specific cells that account for the majority of viral epitope-specific cells in an unseparated memory cell population (Fig 1B). When analyzed for the cell-sorted subsets, frequencies of IE1-and m164-specific IFNγ^+^ functional cells were higher in iTEM (4.8%) and cTEM (1.9 %) compared to TCM (0.7%) (Fig 5A). This makes sense, because viral epitope-specific iTEM and cTEM reflect more recent antigen restimulation in response to antigenic mCMV peptides expressed and presented during the latent infection, whereas cells resulting from past encounters with numerous unrelated antigens quantitatively dominate the TCM pool. Notably, this ranking is just opposite to the ranking in antiviral efficacy. Thus, obviously, the failure of 1,000 iTEM or cTEM in controlling infection (Fig 4B) cannot be explained by a shortage of functional epitope-specific cells.

**Fig 5.**
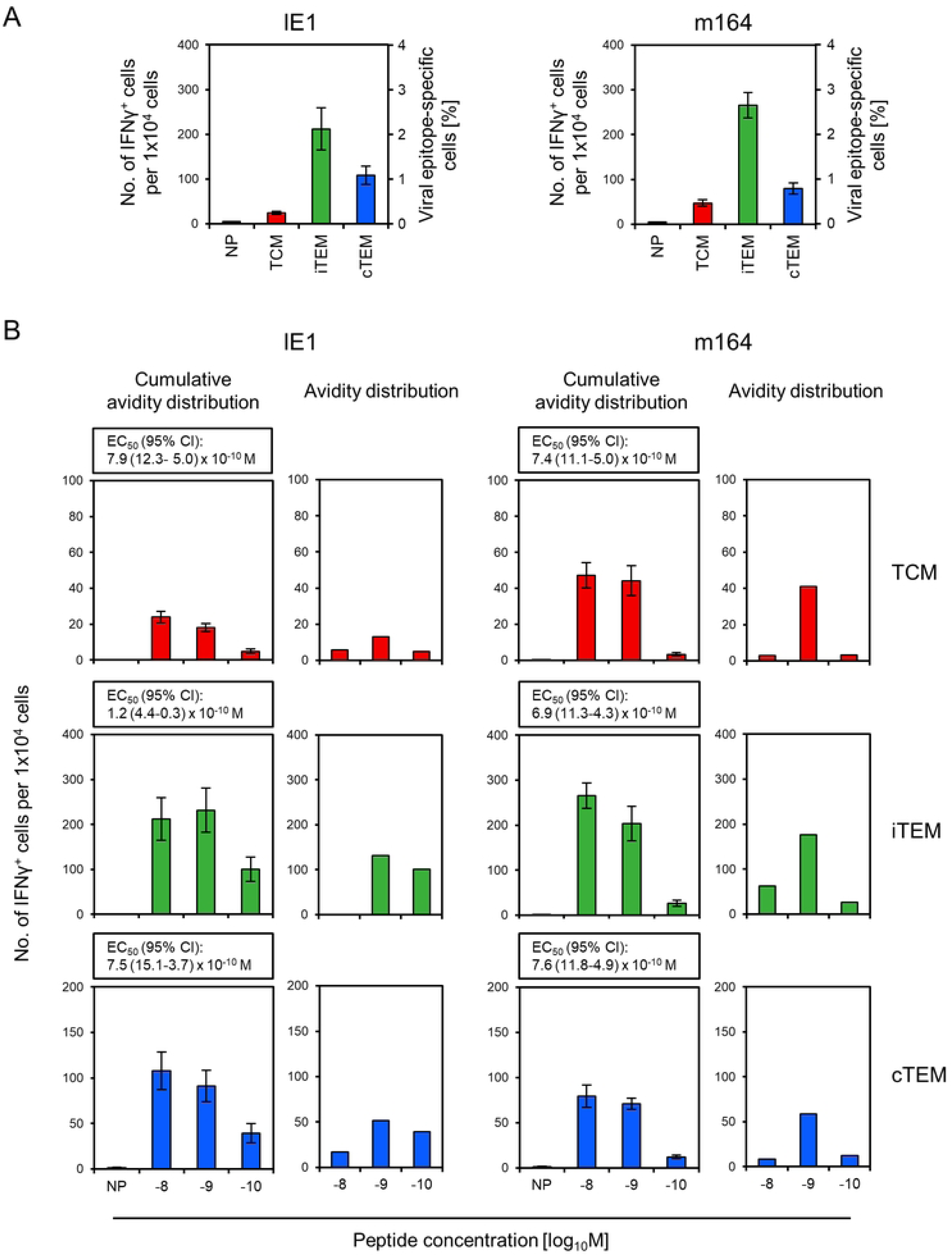
Frequencies and functional avidities of viral epitope-specific memory CD8 T cells differentiated by activation status. (A) Frequencies of viral epitope-specific TCM, iTEM, and cTEM. (Left panel) IE1 peptide-specific cells. (Right panel) m164 peptide-specific cells. Frequencies refer to functional cells responding with secretion of IFNγ in an ELISpot assay to stimulation by P815 cells loaded with the respective antigenic peptide at the saturating concentration of 10^-8^M. (NP) no peptide. (B) Cumulative avidity distributions and deduced Gaussian-like avidity distributions of TCM, iTEM, and cTEM specific for antigenic peptides IE1 and m164, corresponding to (A). Stimulator cells in the ELISpot assay were P815 cells loaded with the respective antigenic peptide in the graded concentrations indicated. (NP) no peptide. (EC_50_ and 95% CI) effective concentration and its 95% confidence interval of antigenic peptide that leads to the half-maximal response of the CD8 T-cell population tested. Throughout, bars represent frequencies determined by intercept-free linear regression analysis. Error bars represent the 95% confidence intervals. Data for transferred CD8 T-cell subsets are color-coded as defined in Fig 4A.

These data also lead to a refined view on the protection data (Fig 4) that had revealed a highly significant protection by AT of 1,000 total TCM. As, in this experiment, IE1- and m164-specific cells together accounted for ∼0,7 % of total TCM, which can be extrapolated to ∼1 % being specific for all viral epitopes, ∼10 transferred viral epitope-specific TCM actually accounted for the observed protection. Note that such numbers show variance and should not be mistaken as being absolutely precise, but they give a decent idea of the scale.

### The poor antiviral activity of iTEM and cTEM is not explained by low TCR avidity

The antiviral efficacy of AT largely depends on the strength and duration of the interaction between peptide-MHC class-I (pMHC-I) complexes presented at the cell surface of infected cells and the cognate TCRs of the CD8 T cells. This determines the intensity of signaling for CD8 T-cell activation and triggering of effector functions. A key parameter is the ‘structural avidity’ measured as the k_off_ rate that quantifies the dissociation of monomeric pMHC-I ligands from the TCRs on living cells [79]. Specifically, the mouse AT model using CTLL selected for high or low ‘functional avidity’ in recognition of the mCMV epitope m164 [49] demonstrated a causal link between high functional avidity, low k_off_ rate, and high protective capacity upon AT [49,79]. Interaction avidity is particularly critical in the case of viruses that express ‘immune evasion’ proteins that interfere with the MHC class-I pathway of antigen presentation (for review, see [20]). As we have shown in the mCMV model, these proteins fail to completely prevent but limit antigen presentation and thereby raise the avidity threshold for CD8 T cells to become sensitized (for review, see [80]). So, only high-avidity CD8 T cells recognize infected host tissue cells and protect upon AT when the virus encodes immune evasion proteins. In contrast, cells infected with an immune evasion gene deletion mutant can be recognized also by low-avidity CD8 T cells, resulting in protection upon AT [49,80].

As our study was performed throughout with mCMV encoding the full set of known immune evasion proteins operating in the MHC class-I pathway [20], protection rested on high-avidity CD8 T cells. The threshold avidity for protection in the presence of immune evasion proteins can be defined by the capacity of CD8 T cells to recognize cells exogenously loaded with synthetic antigenic peptides at loading concentrations of < 10^-9^ M [49,80]. We therefore first tested if unseparated memory CD8 T cells specific for epitopes IE1 and m164 also fulfill this condition, as it was actually predicted by the already demonstrated protective antiviral function [78]. Indeed, the unseparated memory CD8 T-cell population included cells capable of recognizing target cells loaded with either of the two antigenic peptides at a concentration of 10^-10^ M. The half-maximal effective concentration (EC_50_) values, which describe the population average, were also below the protection threshold value of 10^-9^ M (S3 Fig).

To our knowledge, the question of whether memory CD8 T-cell subsets differ in their functional avidities has never been addressed experimentally. We took into consideration that the poor protection by iTEM and cTEM might result from lower functional avidities compared to TCM. This possible explanation was clearly refuted by the data (Fig 5B). Specifically, as required for the control of infection, all three subsets included cells capable of recognizing the two antigenic peptides at a peptide loading concentration of 10^-10^ M, and also the EC_50_ values were below the protection threshold value of 10^-9^ M for all three subsets and for both immunodominant peptides (Fig 5B).

### Only TCM progeny form fully developed NIF, the histological correlate for antiviral protection

From the very low numbers of viral epitope-specific cells required in AT for control of infection in various organs it becomes intuitively clear that protection is not accomplished by an instant effector function of the few initially transferred cells. Instead, protection must result from the effector function of the progeny cell population generated by *in vivo* clonal expansion of the transferred cells [33,78]. As we have shown previously [78], as well as here (Fig 3), AT of unseparated memory cells leads to CD8 T-cell tissue infiltrates that are not randomly distributed but accumulate at infected cells in a micro-anatomical structure, the NIF. By formation of NIF and executing their effector function within the NIF, the infiltrating CD8 T cells confine and eventually resolve tissue infection.

Here we tested the formation of NIF after AT of sorted subsets TCM, iTEM, and cTEM from the experiment shown in Fig 4, all with the same starting cell number of 10^3^ transferred cells (Fig 6). Representative 2C-IHC images of liver tissue sections stained for CD8 and the viral IE1 protein illustrate that fully developed NIF are formed only after AT of TCM, whereas progeny of cTEM form incomplete NIF, and NIF are absent after AT of iTEM (Fig 6A). Quantitation of liver tissue-infiltrating CD8 T cells revealed a rank order of TCM > cTEM > iTEM, anti-correlated to the numbers of infected liver tissue cells in the rank order of TCM < cTEM < iTEM (Fig 6B). A 2-dimensional analysis of NIF size, with the criteria of number of NIF and median number of CD8 T cells per NIF, confirmed the ranking of TCM > cTEM > iTEM (Fig 6C). In conclusion, according to all parameters tested, control of infection is most efficient after AT of TCM, as only TCM progeny infiltrate tissue in numbers needed to form fully developed NIF confining the infection.

**Fig 6.**
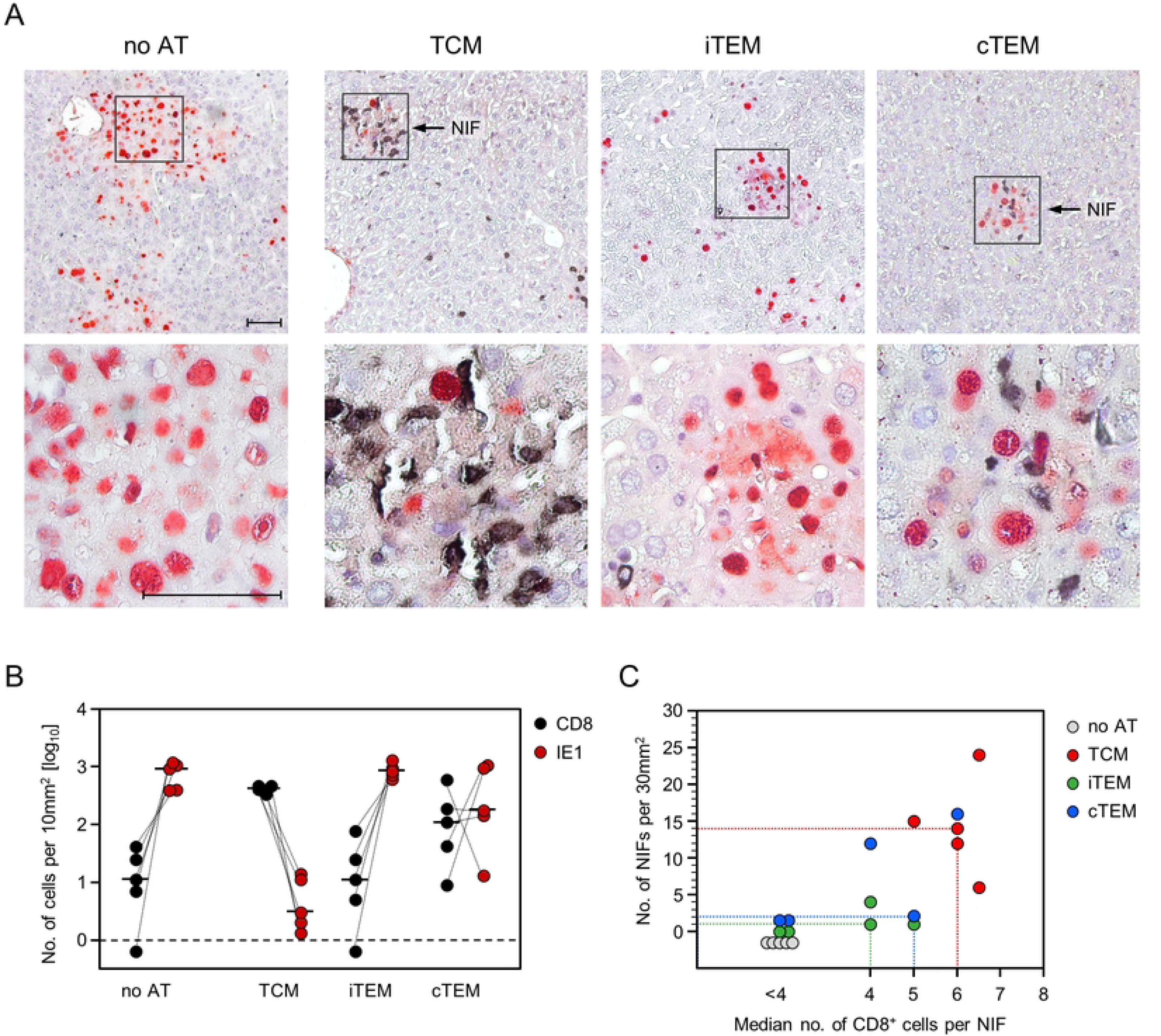
Visualization of liver tissue infection and CD8 T-cell infiltration differentiated by activation status prior to AT. (A) 2C-IHC images of liver tissue sections showing virus control on day 11 by progeny of 10^3^ TCM, iTEM, or cTEM, corresponding to titers of infectious virus shown in Fig 4. Infected liver cells are identified by red staining of the intra-nuclear viral protein IE1. Tissue-infiltrating CD8 T cells are identified by black staining of the CD8a molecule. Light hematoxylin counterstaining reveals the context of liver tissue. (Upper panel images) Low-magnification overviews, showing the confinement of infection by formation of nodular inflammatory foci (NIF) selectively after AT of TCM. Frames demarcate regions of interest resolved to greater detail in the lower panel images. Bar markers represent 50µm. (B) Quantitation of infected IE1^+^ liver cells (red dots) and tissue-infiltrating CD8 T cells (black dots) in representative 10-mm^2^ liver tissue section areas. Symbols represent cell counts for AT recipients tested individually. Linked data are connected by dotted lines, median values are marked. The dashed line indicates the detection limit of the assay. (C) Correlation of the number of NIFs per representative 30-mm^2^ liver tissue section areas with the median number of CD8 cells per NIF, differentiated by transferred CD8 T-cell subset. Symbols represent data from individual mice. The dotted lines mark the AT recipients with the median number of NIFs.

### The superior protection by AT of TCM is explained by their high in situ proliferative potential

The 2C-IHC analysis detecting CD8 T cells and infected cells in their tissue context have revealed the highest numbers of liver tissue-infiltrating CD8 T cells after AT of TCM (Fig 6), although the number of initially transferred cells was the same for all three subsets and even though the number of viral epitope-specific cells was actually lowest in the TCM population (Fig 5A). It was therefore reasonable to propose a higher proliferation rate of TCM compared to cTEM and iTEM.

Standard assays for *in vivo* proliferation are based on loss of a fluorescent reporter dye with every cell division [81,82], but this approach requires AT of high cell numbers, and the resolution is limited to few cell divisions until fluorescence intensity falls below the detection limit. To overcome such technical limitations, we used here our previously described approach of a quantitative ‘input-output’ comparison of the number of initially transferred viral epitope-specific CD8 T cells with the absolute number of progeny cells present in a host organ at defined times after AT [33,78]. CD8 T cells in the stage of cell division were detected *in situ* by 2C-IHC specific for the CD8 molecule, which localizes to the cytoplasm and cell membrane, and the intra-nuclear ‘proliferating cell nuclear antigen’ PCNA (Fig 7A, images). PCNA is a credible marker for currently proliferating cells, as it is expressed during the cell cycle in the G1 phase, reaches its maximum in the S phase and declines during the G2/M phase [83–85]. Compared to *ex vivo* detection of proliferation markers, 2C-IHC has the distinct advantage of localizing proliferating cells in infected tissues, thus visualizing them in their micro-anatomical context [86]. To avoid confusion, it must be noted that productively infected cells also express PCNA in the nucleus, reflecting viral DNA replication activity [87]. This poses no problem, as PCNA^+^ infected liver cells do not co-express CD8.

**Fig 7.**
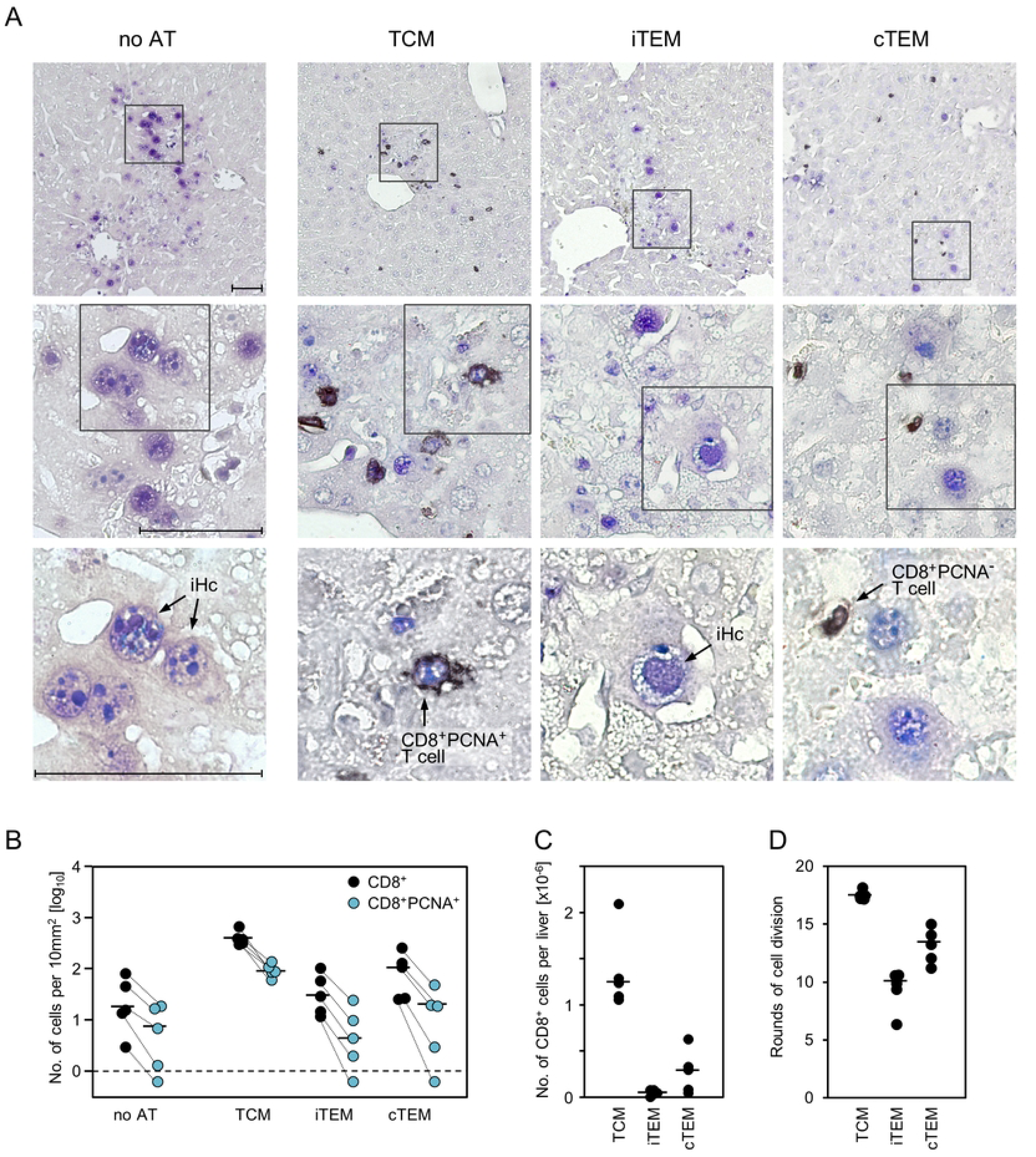
Visualization of extra-lymphoid proliferation and whole-organ quantitation of memory CD8 T-cell progeny differentiated by the activation status prior to AT. (A) 2C-IHC images of liver tissue sections illustrating *in situ* proliferating CD8 T cells after transfer of 10^3^ TCM, iTEM, or cTEM. (no AT) control group with no AT performed. Liver tissue-infiltrating CD8 T cells are identified by black staining of the CD8a molecule. The ‘proliferating cell nuclear antigen (PCNA)’ is identified by blue nuclear staining. (CD8^+^PCNA^+^ T cell) Proliferating CD8 T cells are identified by co-expression of intranuclear PCNA, stained in blue, and cytoplasmic as well as membrane CD8a, stained in black. (CD8^+^PCNA^-^ T cell) CD8 T cells not proliferating at the time of analysis. (iHc) productively infected liver cells, most of which are hepatocytes, also express PCNA in their nuclei. Light hematoxylin counterstaining reveals the context of liver tissue. (Upper panel images) Low-magnification overviews. Frames demarcate regions of interest resolved to increasingly greater detail in the center and lower panel images. Arrows point to representative cells of interest. Bar markers represent 50µm throughout. (B) Quantitation of all liver tissue-infiltrating CD8^+^ cells (black dots) and of proliferating CD8^+^PCNA^+^ cells as a subset thereof (blue dots) in representative 10--mm^2^ liver tissue section areas. Symbols represent cell counts for AT recipients tested individually. Linked data are connected by dotted lines, median values are marked. The dashed line indicates the detection limit of the assay. (C) The absolute numbers of CD8 T cells, present in the whole liver of the AT recipients on read-out day 11, were calculated by extrapolation based on the numbers counted in the tissue sections. Symbols represent data from mice tested individually, with the median values marked. (D) The numbers of CD8 T-cell divisions that have occurred until the read-out day 11 were calculated based on the numbers of viral epitope-specific cells present in initially transferred 10^3^ cells of the memory CD8 T-cell subsets TCM, iTEM, and cTEM. Symbols represent data for individual AT recipients. Median values are marked.

In accordance with the 2C-IHC analysis of CD8 T cells and IE1-expressing infected liver cells shown above (Fig 6), the number of liver-infiltrating CD8 T cells ranked TCM > cTEM > iTEM, and this ranking also applied to proliferating CD8^+^PCNA^+^ cells (Fig 7B). Throughout, CD8^+^PCNA^+^ cells present at the time of analysis represented only a fraction of the tissue-infiltrate CD8 T cells. This is easy to explain, as PCNA^-^ tissue-infiltrate CD8 T cells likely have completed proliferation and were thus back to the cell-cycle G0-phase at the time of analysis. Absolute cell counts, extrapolated from tissue sections to the total liver (for the formula, see [33]), revealed an almost absence of iTEM progeny and a strong predominance of TCM progeny over cTEM progeny (Fig 7C), corresponding to integer median cell division numbers (ranges) of 17 (17-18) for TCM, 10 (6-11) for iTEM, and 13 (11-15) for cTEM (Fig 7D).

## Discussion

In clinical immunotherapy to prevent CMV disease in HCT recipients, providing sufficient cell numbers for AT remains a logistical challenge. Early clinical protocols followed the strategy to expand low numbers of memory CD8 T cells derived from a ‘CMV-antibody seropositive’ donor to high cell numbers in cell culture, that is, by establishing clonal or polyclonal cytolytic T-lymphocyte lines (CTLL) [34,35,51]. Translated from the language of clinicians to the language of viral immunologists, the presence of CMV-specific antibodies is merely an indicator for a past primary CMV infection that has developed into a latent infection, characterized by absence of infectious virus, maintenance of the viral genome in certain cell types, and presence of CMV-specific immunological memory (reviewed in [88], and recently updated for mCMV in [63,64]).

In the early days, clinical AT focused entirely on CTLL specific for the abundant hCMV tegument protein pp65/UL83, as the use of virions for the restimulation in cell culture had led to selection to the favor of pp65/UL83-specific CTLL [89–92]. Unbiased determination of the specificity spectrum of the human memory CD8 T-cell response to hCMV by viral genome-wide antigenicity screening confirmed pp65/UL83 as a significant source of antigenic peptides, but also revealed an antigenicity of most hCMV ORFs at broad HLA coverage, and still of several ORFs for the HLA set of any individual [93]. Accordingly, for any HLA class-I matched donor-recipient pair, CD8 T-cells of several specificities qualify as candidates for use in AT.

An important contribution of the mouse model was the finding that expansion of memory CD8 T cells to high numbers of effector cells in cell culture comes at the cost of a loss of protective activity per cell, thus negating the effort to establish CTLL [47,48]. Clinical data also revealed a superior protection capacity of *ex vivo* isolated memory CD8 T cells [37,38,50] when compared to published experience with CTLL, but only the mouse model allowed the ‘Proof of Concept’ by a direct comparison between *ex vivo* sort-purified memory CD8 T cells and CTLL, specific for the same viral epitope and tested in parallel in the same experiment. This comparison revealed more than two orders of magnitude higher protective capacity of the memory cells [47,48].

While the use of memory CD8 T cells for clinical AT is meanwhile standard, less attention is paid to potentially different properties of memory cell activation and differentiation subsets, specifically of central memory cells (TCM) and effector-memory cells (TEM), which can be subdivided into conventional TEM (cTEM) and inflationary TEM (iTEM). ‘Stemness’ with high self-renewal capacity associated with a high proliferation rate is a reported advantage of CD62L^+^ TCM compared to CD62L^-^ TEM in mouse models of bacterial pathogen-specific AT [38,94], and, in analogy, low-dose clinical AT of hCMV-specific human CD8^+^ T cells into HCT patients led to a vigorous expansion of the transferred cells [38]. The clinical hCMV part of the study by Stemberger and colleagues [38], however, depended on compassionate-use settings that did not allow a distinction between TCM and TEM. Nonetheless, based on these studies, one can take it for granted that TCM generally have a superior proliferation capacity. On the other hand, a protective effect of TCM comes with delay, whereas, as the name ‘effector-memory cell’ indicates, TEM might have the advantage of a more rapid antiviral effect.

In latent CMV infections, the viral genomes are not completely silenced. Instead, episodes of viral gene desilencing lead to limited and transient transcription that does not follow the coordinated gene expression cascade of the productive viral cycle [95–98]. Linking this to immune surveillance of latent infection, the murine model has shown stochastic and transient expression also of viral genes that code for antigenic peptides [60,61]. Their presentation on the surface of latently infected cells, meanwhile identified as endothelial cell types, is proposed to drive memory inflation (MI) [55,61,62]. The stochastic nature of the transcription episodes is perfectly reflected by stochastic clonal expansions of virus epitope-specific CD8 T cells [99] and explains the variance in viral epitope-specific CD8 T-cell reconstitution dynamics between different HCTs performed with an identical protocol [24].

MI is defined by expansion of the pool of KLRG1^+^CD62L^-^ TEM, previously referred to as ‘short-lived effector cells (SLEC)’ [53], which we have proposed to re-name inflationary TEM (iTEM) in distinction from non-inflationary KLRG1^-^CD62L^-^ conventional TEM (cTEM) [52]. MI is favored by conditions of primary infection that lead to a high load of latent viral DNA, since a high number of latent viral genomes as templates obviously increases the probability for episodes of antigen-coding transcription. One parameter, experimentally shown to favor MI in the immunocompetent mouse model, is the initial dose of infection [100,101]. However, regardless of the initial dose of infection, which is probably always very low in humans, subsequent virus spread in the host that largely depends on the immune status at the time of infection and which is high in the immunologically immature or immunocompromised host, is likely even more important to establish a high latent viral genome load favoring MI [100,102]. Dependence on the individual infection history, defining the magnitude of productive infection and thus also the latent viral DNA load, explains why MI is not consistently observed in humans ([103], for a commentary, see [104]). The human counterpart of iTEM, that is, the cells expanding during MI and thus defining MI, show an ‘advanced differentiation phenotype’ that also includes high expression of KLRG1 and low expression of CD62L [105]. In both, human and murine latent CMV infections, the inflationary CD8 T cells are functional (reviewed in [106]). Specifically, in mCMV, MI during viral latency was originally described as expansion of CD62L^-^ cells capable of secreting IFNγ upon stimulation by antigenic peptide presentation [107], and SLEC/iTEM were found not to be terminally differentiated effector cells but to proliferate upon AT [53]. Based on their high numbers under conditions of MI, facilitating their isolation, and on their functionality and proliferative potential, iTEM were promising candidates for controlling CMV infection. As far as we are aware of, although iTEM were proposed to surveil viral latency by sensing and terminating productive viral reactivation [60], their propensity to control acute infection was never tested in an AT model.

Controlled studies comparing the protective efficacies of memory CD8 T-cell subsets in parallel by AT into recipient hosts are not feasible in humans, because individual HCT patients differ in their genetics and progress of hematopoietic reconstitution, in latent virus strain(s)/clinical isolates that can differ in host-cell tropism [1,108–113], in latent viral genome load depending on the individual history of infection, in the time of virus reactivation onset and organ site of reactivation, and in the extent of inter-as well as intra-tissue virus spread. In addition, there is no way to find identical HCT and AT donors, who also differ in their memory-shaping infection history, including AT donor latent virus strain and epitope-specificity composition of the memory cell pool. Every single one of these unavoidable variables in both AT donor and recipient renders a quantitative comparison between memory CD8 T-cell subsets by clinical AT a lottery game. It is this type of question that can reliably be answered only in an animal model [21].

And the answer was clear: under conditions of low-dose AT, performed in absence of HCT to reduce the number of variables, only TCM were able to clear productive infection by infiltrating infected tissue in numbers high enough to form NIF, the micro-anatomical correlate of protection. All three memory CD8 T-cell subsets were able to control infection after high-dose AT. Protection by cTEM, and in particular by iTEM at high doses may indeed reflect their more rapidly exerted antiviral effector function, but this is of little help for clinical AT where the provision of high cell numbers is the logistically limiting factor for AT. Notably, the number of iTEM required for control of the infection is strikingly similar to the experience made previously with CTLL [47,48]. In fact, iTEM and the majority of the cells of a CTLL share the ‘advanced differentiation phenotype’ of high KLRG1 and low CD62L expression. Notably, as shown previously for a short-term CD8^+^ CTLL specific for the IE1 peptide and propagated by repetitive restimulation in cell culture, the CTLL population split into a majority of iTEM-like KLRG1^+^CD62L^-^ cells and a minority of cTEM-like KLRG1^-^CD62L^-^cells [49]. These are precisely the two phenotypes that are of low efficacy in AT, with iTEM < cTEM [this report]. We thus would like to put forward the interpretation that CD8 T cells repetitively restimulated by stochastically expressed and presented antigenic peptides in the latently infected host in a sense resemble CTLL, with the difference that CTLL also contain terminally-differentiated, cytolytic TEC, whereas *ex vivo* isolated viral epitope-specific CD8 T cells are cytolytically active only in the acute phase of infection but no longer during latent infection [114].

Attempting to define the reason for the higher protective activity of TCM in low-dose AT compared to iTEM and cTEM, we have excluded a higher frequency of viral epitope-specific cells or a higher functional avidity in recognizing antigenic peptides presented on infected cells. The most distinctive difference between TCM and both subsets of TEM was the superior viral epitope-specific infiltration into infected host tissues and the formation of NIF, as shown here exemplarily for the liver. This cannot be explained by a more efficient homing of the initially transferred CD62L^+^ TCM to non-lymphoid tissues, because CD62L, also known as L-selectin, is a ‘homing receptor’ expressed by recirculating cells for mediating their temporal homing to lymphoid tissues. It appears that TCM must first be activated by antigen encounter to convert to CD62L^-^ TEM for infiltrating non-lymphoid tissues [115,116]. In accordance with this, as we have shown recently for the location of viral epitope-specific CD8 T cells within latently infected lungs [117], CD62L^-^ TEM efficiently migrate from the intravascular compartment (IVC) to the extravascular compartment (EVC), the lung parenchyma, where in particular cTEM were found enriched. In contrast, CD62L^+^ TCM are absent in the EVC, which is best explained by loss of CD62L and conversion to TEM during transmigration of the lung endothelium. From all this we conclude that TEM exogenously administered to the IVC by AT will also efficiently infiltrate infected host tissues.

So, what remains as an explanation for the much better antiviral control by AT of TCM is their superior proliferation capacity that leads to a massive expansion of the pool of protective TEC. It is a strength of our approach that we visualized intra-tissue virus spread and its prevention by infiltrating, NIF-forming CD8 T cells in 2C-IHC. Notably, using PCNA as a credible marker for proliferation, we could demonstrate an *in situ* proliferation of CD8 T cells at a non-lymphoid site of viral pathogenesis. An ‘input-output’ quantitation of the progeny of the transferred cells allowed us to calculate the number of cell cycles passed after AT. It should be noted that these data represent minimum estimates based on cells present in the liver, although it is clear that progeny of transferred cells control the infection also in other target organs of CMV disease. Accordingly, the true number of cell cycles must be corrected upwards. On the other hand, for the sake of ease, we have neglected proliferation of CD8 T cells specific for subdominant viral peptides, so that the true number of cell cycles must be corrected downwards. These cells however, collectively account for < 50% of the epitope-specific cells. Thus, the two corrections are in opposite directions and certainly do not interfere with the relevant message of ranking the proliferation capacity in the order of TCM >> cTEM > iTEM. It also must be considered that every factor of 2 in correcting the number of initially transferred cells reaching the liver results in a difference of just one cell cycle.

Although the absolute number of iTEM progeny in the liver was almost negligible, our data are not conflicting with the finding by Snyder and colleagues [53], who have shown that fluorescence-labeled iTEM proliferate upon AT. We have estimated 10 (6-11) cell cycles for iTEM and 17 (17-18) cell cycles for TCM. This does not sound like a dramatic difference, unless one has successfully attended a math lecture on exponential functions. High absolute numbers are achieved by the late cell divisions, while the early cell divisions make only a small contribution.

In conclusion, low-dose AT of iTEM or cTEM in HCT patients is no promising option, as protection against CMV disease depends on vigorous expansion of virus-specific TCM.

## Materials and methods

### Ethics statement

Animal experiments were performed in accordance with the national animal protection law (Tierschutzgesetz (TierSchG)), animal experiment regulations (Tierschutz-Versuchstierverordnung (TierSchVersV)), and the recommendations of the Federation of European Laboratory Animal Science Association (FELASA). The experiments were approved by the ethics committee of the Landesuntersuchungsamt Rheinland-Pfalz, permission numbers 177-07/G14-1-015 and 177-07/G19-1-049.

### Mice, viruses, and route of infection

Female BALB/cJ (haplotype H-2^d^) mice were bred and housed under specified-pathogen-free (SPF) conditions by the Translational Animal Research Center (TARC) at the University Medical Center of the Johannes Gutenberg-University Mainz. Immunocompetent CD8 T-cell donors and immunocompromised recipients were used at an age of 8-to-12 weeks.

mCMV (strain Smith, ATCC VR-1399) was used as wild-type virus (mCMV-WT). BAC-derived recombinant viruses mCMV-IE1-L176A and the corresponding revertant mCMV-IE1-A176L were described previously [60]. All viruses were propagated in cell culture and purified by standard methods [118]. Intraplantar infection was performed by injection of 10^5^ plaque-forming units (PFU) of the respective virus into the left hind footpad.

### Immunnomagnetic enrichment of memory CD8 T cells and fluorescence-based cell sorting

Immunocompetent BALB/cJ mice were infected with mCMV-WT for use as T-cell donors. After 8 months, CD8 T cells derived from a pool of 10-12 spleens were enriched either by immunomagnetic positive selection [119], or, in cases of subsequent fluorescence-based cell sorting, by immunomagnetic negative selection using the MagniSort^TM^ mouse CD8 T-cell enrichment kit (catalog no. 8804-6822-74; eBioscience), following the manufacturer’s instructions.

Memory CD8 T-cell subsets were isolated by fluorescence-based cell sorting after staining of CD8 and activation markers CD62L and KLRG1, using FITC-conjugated anti-CD8a (clone 53-6.7; eBioscience), PE/Dazzle594-conjugated anti-KLRG1 (clone 2F1; BioLegend), and PE-Cy7-conjugated anti-CD62L (clone MEL-14; eBioscience). Sort gates were set on CD8 T cells and CD8 T-cell subsets TCM (CD62L^+^KLRG1^-^), iTEM (CD62L^-^KLRG1^+^), and cTEM (CD62L^-^KLRG1^-^). IE1-specific memory CD8 T cells were purified by fluorescence-based cell sorting using PE-conjugated, IE1 peptide-folded MHC-I dextramers H-2Ld/YPHFMPTNL (Immudex, Copenhagen, Denmark). Analyses and cell sorting were performed with flow cytometer BD FACSAria^TM^ I and FACSDiva analysis software (BD Biosciences).

### Cytofluorometric (CFM) analyses

Enrichment of CD8 T cells by immunomagnetic selection was documented by CFM quantitation of TCRβ^+^CD8^+^CD4^-^ lymphocytes. Unspecific staining was blocked with unconjugated anti-FcγRII/III antibody (anti-CD16/CD32, clone 93; BioLegend). Specific staining for 3-color CFM analysis was performed with PE-conjugated anti-TCRβ (clone H57-597; BD Bioscience), PE-Cy5-conjugated anti-CD8a (clone 53-6.7; eBioscience), and FITC-conjugated anti-CD4 (clone GK1.5; BioLegend) antibodies. For distinguishing antigen-experienced memory CD8 T-cell subsets from naïve CD8 T cells, a 4-color CFM analysis was performed with PE-Cy7-conjugated anti-CD44 (clone IM7; eBioscience), PE/Dazzle594-conjugated anti-CD8a (clone 53-6.7; BioLegend), FITC-conjugated anti-KLRG1 (clone 2F1; eBioscience), and PE-conjugated anti-CD62L (clone MEL-14; eBioscience) antibodies. All analyses were performed with flow cytometer Cytomics FC500 and CXP analysis software (Beckman Coulter).

### Adoptive transfer (AT) of memory CD8 T cells

For AT of donor-derived memory CD8 T cells or subsets thereof, 8-week-old BALB/cJ recipients were immunocompromised by hematoablative total-body γ-irradiation with a single dose of 6.5 Gy. Donor cells were transferred 4 hours later by intravenous infusion. Recipient mice were infected 2 hours after AT with mCMV-WT. In the specific case of AT of IE1 epitope-specific memory CD8 T cells, recipients were infected either with mutant virus mCMV-IE1-L176A or revertant virus mCMV-IE1-A176L. Throughout, mice left without AT served as controls for unrestricted viral replication.

### Peptides and quantitation of functional epitope-specific memory CD8 T cells

Viral epitopes corresponding to antigenic peptides presented by MHC class-I molecules K^d^, D^d^, and L^d^ are derived from the mCMV ORFs m18, M83, M84, M105, m123/IE1, m145, and m164 (amino acid sequences and presenting MHC class-I molecules listed in [49]). Custom peptide synthesis with a purity of > 80% was performed by JPT Peptide Technologies.

Immunomagnetically-purified CD8 T cells and sorted subsets thereof, derived from pooled spleens of AT donor mice latently infected with mCMV-WT, served as responder cells in IFNγ-based enzyme-linked immunospot (ELISpot) assays ([78], and references therein). In essence, for quantitating functional, mCMV epitope-specific memory CD8 T cells, the corresponding synthetic peptides were exogenously loaded on P815 (H-2^d^) mastocytoma cells at the molar concentrations indicated, for serving as stimulator cells in the assay. Graded numbers of responder cells were seeded with the peptide-loaded stimulator cells in triplicate microcultures. After 18 hrs of incubation, spots were counted using the ImmunoSpot S4 Pro Analyzer (Cellular Technology Limited).

### ELISpot assay calculations and determination of avidity distributions

Frequencies (most probable numbers, MPN) of cells responding in the ELISpot assay and the corresponding 95% confidence intervals were calculated by intercept-free linear regression analysis from the linear portions of regression lines based on spot counts from triplicate assay cultures for each of the graded cell numbers seeded [120]. Calculations were performed with SPSS Statistics, Version 23.

Cumulative avidity plots show the measured frequencies of cells responding at the indicated peptide concentration and all lower concentrations. Based on MPN and the corresponding upper and lower 95% confidence limits, half-maximal effective concentration (EC_50_) values, representing peptide concentrations that result in the half-maximal response of the cell population, were calculated with Quest Graph™ EC_50_ Calculator (AAT Bioquest, Inc.; retrieved from https://www.aatbio.com/tools/ec50-calculator). Gaussian-like avidity distributions reveal frequencies of cells with an avidity defined precisely by the peptide concentration indicated. These are deduced from the cumulative avidity distribution values by plotting the response increments between a certain peptide concentration and the next lower peptide concentration [52].

### Visualization of tissue infection, CD8 T-cell infiltration, and CD8 T-cell proliferation

At 11 days after AT, infectious virus in spleen, lungs, and liver was quantitated in whole organ homogenates by a virus plaque assay performed on monolayers of mouse embryo fibroblasts under conditions of ‘centrifugal enhancement of infectivity’ ([119], and references therein).

To visualize and quantitate infection and CD8 T-cell infiltration in the micro-anatomical context of liver tissue, infected cells and CD8 T cells were identified in tissue sections by 2C-IHC staining of the intra-nuclear viral immediate-early (IE) protein IE1 in red color and the CD8a molecule in black [78]. To detect *in situ* proliferating CD8 T cells, 2C-IHC was used for visualizing cells co-expressing the CD8a molecule (black staining, see above) and PCNA (blue staining). Blue staining of PCNA was achieved by using a species-cross reactive mouse IgG2a-kappa monoclonal antibody directed against PCNA (clone PC10; BD Bioscience) and the Vector® Blue Substrate Kit, Alkaline Phosphatase (AP) (catalog no. SK-5300; Vector Laboratories).

### Quantitation of infected tissue cells and tissue-infiltrating CD8 T cells

Total numbers of infected cells and of tissue-infiltrating CD8 T cells were calculated from cells counted in representative tissue sections and extrapolated to the whole organ by using a mathematical formula that corrects for overestimation when the diameter *D* of the counted object is > the thickness *d* of the tissue section (for a detailed explanation, see [33]).

### Calculation of the number of cell divisions

The number of cell divisions was calculated as *n* = log_2_ [*N(t)/N(0)*], where *N(t)* = total number of tissue-infiltrate CD8 T cells per whole organ at time *t* after AT and *N(0)* = number of initially (*t* = 0) transferred viral epitope-specific cells of CD8 T-cell subsets TCM, iTEM, and cTEM.

### Statistical Analysis

To evaluate statistical significance of differences between two independent sets of data, the two-sided unpaired t-test with Welch’s correction of unequal variances was used. Differences were considered statistically significant for P-values (*) <0.05, (**) <0.01, and (***) <0.001. Calculations were performed with Graph Pad Prism 6.04 (Graph Pad Software, San Diego, CA).

## Acknowledgments

The authors thank Sebastian Attig, Department of Translational Oncology and Immunology at the Institute of Immunology, University Medical Center of the Johannes Gutenberg University Mainz, Mainz, Germany, for CD8 T-cell sorting.

## Author Contributions

**Conceptualization:** Matthias J. Reddehase. Rafaela Holtappels, Niels A. Lemmermann

**Data Curation:** Rafaela Holtappels, Niels A. Lemmermann, Jürgen Podlech, Sara Becker

**Formal analysis:** Rafaela Holtappels, Niels A. Lemmermann, Sara Becker, Sara Hamdan. Kirsten Freitag

**Funding acquisition:** Rafaela Holtappels, Niels A. Lemmermann, Matthias J. Reddehase

**Investigation:** Sara Hamdan, Sara Becker, Kirsten Freitag, Jürgen Podlech

**Methodology:** Rafaela Holtappels, Sara Becker, Sara Hamdan, Kirsten Freitag, Jürgen Podlech

**Project administration:** Rafaela Holtappels, Niels A. Lemmermann, Matthias J. Reddehase

**Supervision:** Rafaela Holtappels, Niels A. Lemmermann, Jürgen Podlech

**Validation:** Rafaela Holtappels, Niels A. Lemmermann, Matthias J. Reddehase.

**Visualization:** Rafaela Holtappels, Sara Becker, Jürgen Podlech, Kirsten Freitag.

**Writing – Original Draft Preparation:** Matthias J. Reddehase

**Writing – Review & editing:** Rafaela Holtappels, Niels A. Lemmermann

## Data Availability Statement

All relevant data are within the manuscript and its Supporting Information files.

## Supporting information

**S1 Fig. Activation phenotypes of CD8 T cells.** Comparative CFM analyses of cell surface marker expression by CD8 T cells derived from the spleen of age-matched uninfected BALB/c mice (upper panels) and of memory CD8 T cells derived from the spleen of BALB/c mice in the stage of latent infection (lower panels). Shown are color-coded 2D fluorescence density plots for the cell surface marker combinations indicated, with red and blue color representing highest and lowest cell numbers, respectively. Gates were set on CD8 T cells in the SSC (sideward scatter) versus CD8 plots.

**S2 Fig. Control of infection by AT of total memory CD8 T cells and activation subsets: pilot experiment. (A)** AT of unseparated memory CD8 T cells. For details, see the Legend of Fig. 2 in the body of the text. (B) AT of memory CD8 T-cell subsets TCM, iTEM and cTEM. For details, see the Legend of Fig. 4. Asterisk-coded statistical significance levels for differences between the AT groups and the no-AT control group (Ø): P < (**) 0.01, and (***) 0.001. (n.s.) not significant.

**S3 Fig. Frequencies and functional avidities of viral epitope specific memory CD8 T cells.** Shown are cumulative avidity distributions and deduced Gaussian-like avidity distributions of unseparated memory CD8 T cells specific for antigenic peptides IE1 and m164. For details, see the Legend of Fig. 5 in the body of the text.

